# The role of relA-mediated stringent response on the nutritional, environmental, antimicrobial resistance and biofilm formation in Klebsiella pneumoniae and its effect on Drosophila melanogaster survival

**DOI:** 10.64898/2026.01.04.697604

**Authors:** Rochell T. Davis, Paul D. Brown

## Abstract

The stringent response has far-reaching consequences, with links to antimicrobial resistance, stress and virulence. This study assessed the role of *relA* in nutritional, environmental and antimicrobial stress in *Klebsiella pneumoniae*, the extent of polysaccharide capsule and biofilm formation, and the pathogenic effect on *Drosophila melanogaster*. Two single mutants (KP03*DrelA* and KPPR1*DrelA*) obtained using the Lambda Red Recombinase Technique were the focus of analyses. We assessed environmental (ethanol, osmotic, heat) and nutritional stress (carbon, phosphate, amino acid) tolerance, capsule formation, and cell size determination in wild-type (WT) and mutant bacteria. Biofilm and agar plate susceptibility assays were performed on both starved and non-starved strains using gentamicin and ceftazidime, and stress response genes were analyzed. *Drosophila melanogaster* was used to examine the pathogenic effect of the presence or deletion of *relA* on fly survival. *DrelA* mutants had reduced fitness to environmental and nutritional stress compared to WT strains. Mutant strains’ cell lengths were elongated and lacked a capsule versus WT and non-starved strains, and mutant strains exhibited enhanced biofilm formation or survival in the presence of ceftazidime and gentamicin. Stress response genes *rpoD, phoR, phoU* and *pstB* were absent during starvation, but *rpoS* was detected in C^-^, C^+^PO^+^ and serine hydroxamate (SHX) media; *mrkA* gene was not detected during starvation. Lastly, the animal model proved to be effective in showing infection levels associated with the presence of *the relA* gene.

## Introduction

*Klebsiella pneumoniae* is a Gram-negative bacterium from the *Enterobacteriaceae* family. The bacteria are encapsulated, non-motile and ubiquitously found in nature [1, 2, 3]. The bacterium is often considered an opportunistic pathogen with no evidence of pathology, but it can migrate to other tissues thus resulting in life-threatening infections of pneumoniae, urinary tract infections (UTIs), bloodstream infections and sepsis [4]. According to the European Centre for Disease Prevention and Control (EARS-net) [5] more than two thirds of *K. pneumoniae* isolates reported were resistant to at least one antimicrobial group, which includes a combined resistance to fluoroquinolones, third generation cephalosporins and aminoglycosides being the most common resistance phenotypes.

There are four major virulence factors identified in *K. pneumoniae*, namely, lipopolysaccharide (LPS), siderophores, capsule and fimbriae (pilli) [6]. *K. pneumoniae* produces two types of fimbriae on its surface, type 1 used by many enterobacterial species and is mannose-binding specific [7] and type 3 commonly identified in all *K. pneumoniae* species which facilitates adherence to almost all cell types in-vitro [8, 9]. In addition, the type 3 fimbria plays an important role in biofilm formation on abiotic and biotic surfaces but its role in biofilm formation on urinary catheters are yet to be elucidated [10]. Hence, *K. pneumoniae* strains capable of producing biofilm usually result in life threatening infections with increased treatment failure and acquisition of antibiotic resistance mechanisms [11]. Also, other virulence factors have been identified but they have not been thoroughly characterized in terms of clinical significance and mechanisms of action. These other factors are outer membrane proteins (OMPs), porins, efflux pumps, iron transport systems and genes involved in allantoin metabolism [4].

The stringent response (SR) is a broad, highly conserved bacterial stress response ([12]. It is induced by different starvation and stress signals which ultimately has downstream effects on a bacteria’s survival [13, 14], since the bacteria utilizes the response to determine resource allocation for cell maintenance function or growth [12]. The main signal molecule of the SR are the two secondary messenger nucleotide molecules of guanosine 5’-diphosphate-3’-diphosphate (ppGpp) and guanosine 3’-diphosphate 5’- triphosphate (pppGpp) collectively called (p)ppGpp alarmones [15]. Levels of ppGpp are under control by two enzymes produced by two different genes: *relA,* a monofunctional gene, first identified in *E. coli* and since then has been found in most gram-negative bacteria of the Gamma Proteobacteria class*. RelA* synthesizes ppGpp under amino acid starvation *and spoT*, a bifunctional gene has both synthetic and hydrolytic properties to nutrient starvation and numerous other stresses [16]. Furthermore, the accumulation of ppGpp in the cells also results in other physiological functions including bacterial competence [17], thermotolerance [18], and sensitivity to antibiotics [19] are only a few of such effects.

While the effect of *relA* has been studied in numerous bacteria, no work or there is a gap in knowledge in *Klebsiella pneumoniae* in relation to the stringent response and the effect it has on the bacteria’s capacity to withstand periods of stress. Therefore, this study investigated the relationship or contribution of the stringent response to stress tolerance, antimicrobial resistance, virulence, ability to form biofilms and subsequently its implication in pathogenicity.

## Materials and methods

### Bacterial strains and growth conditions

Table 1 contains the bacterial strains and plasmids used to conduct this study. For routine growth of K. *pneumoniae* ATCC 43816 KPPR1 and *K. pneumoniae* wild-type (WT) strains (KP03, KP04, KP18, KP24) Luria Bertani (LB) broth was used and supplemented with the appropriate antimicrobial agents where required. Wild-type strains mentioned above were chosen based on previously unpublished data of antibiotic susceptibility, biofilm formation and ERIC PCR banding pattern [20]. Nutritional starvation and environmental stress were carried out in minimal M9 medium which contained the following reagents (per litre): Na_2_HPO_4_, 6 g; KH_2_PO_4_, 3 g; NaCl, 0.5 g; NH_4_Cl, 1 g; MgSO_4_.7H_2_O, 0.246 g; CaCl_2_.2H_2_O, 14.7 mg with either of carbon (I), phosphate (II) or both nutrients eliminated (III) based on starvation criteria. Also, amino acid starvation (IV) used the serine hydroxamate (SHX) (0.3 mg/ml), an analogue of L-serine. The following antimicrobial agents were used when required: 100 µg/ml ampicillin (Amp), 30 µg/ml rifampicin (Rif), 50 µg/ml ceftazidime (CAZ), 50 µg/ml gentamicin (GEN) and 50 µg/ml kanamycin (Kan) [21, 22].

**Table 1.** Bacterial strains and plasmids used in this study.

| Strains/Plasmids | Description | Reference or source |
| --- | --- | --- |
| ***K. pneumoniae* strains** |  |  |
| ATCC 43816 (KPPR1) | Control strain | ATCC |
| *ΔrelA* mutant of KPPR1 | *relA*::Kan; Kan<sup>r</sup> insertional mutant of *relA* of KPPR1 | This study |
| KP03 | Clinical type; isolated from sputum of patient | CML |
| *ΔrelA* mutant of KP03 | *relA*::Kan; K; Kan<sup>r</sup> insertional mutant of *relA* of KP03 | This study |
| KP04, KP18, KP24 | Clinical type; isolated from sputum of patient | CML |
| **Plasmids** |  |  |
| pKD46 | BW25113 *E. coli* strain; Ap<sup>r</sup> araBp gam bet exo oriR101 | EGSC |
| pkD4 | BW25141 *E. coli* strain Ap<sup>r</sup> Kan<sup>r</sup> oriR6Kgamma rgnB | EGSC |
| pGEM-T vector | Ap<sup>r</sup>, *lacZ* | Promega |

*Note*; CML, Central Medical Laboratory, Jamaica; EGSC, *E. coli* Genetic Stock Center; Kan, kanamycin; Kan<sup>r</sup>, Kanamycin resistant; Ap<sup>r</sup>, Ampicillin resistant; ATCC, American Type Culture Collection

### Detection of a portion of *relA* gene in wild-type (WT) strains

*K. pneumoniae* chromosomal DNA was extracted using kit provide by Promega Genomic DNA Purification. The PCR analysis of *relA* gene was carried out using a 50 µl PCR mixture with primers provided by Integrated DNA Technology (IDT), Coralville, Iowa, USA: *relA* (F)-AGATTGCAAGCATTACACGTC and *relA* (R)- GCATGCAAGCTTGGCACTGGC. (2.5 µl of 10 µM stock solutions), with 25 µl 2x Promega Go*Taq* master mix reaction buffer, containing 400 µM of dNTPs, 3 mM MgCl_2_, 1 U Go *Taq* DNA polymerase (Promega, Madison, WI, USA), 18.5 µl nuclease free water and 2.5 µl of DNA. A thirty-five (35) cycles amplification steps were done consisting of denaturation step at 94°C for 5 min, 54°C for 5 min, 72°C for 2 min (extension) and a final extension step at 72°C for 7 min [23]. A 1.5% agarose gel was used for amplicons already stained in ethidium bromide and were subsequently visualized using 3UV Benchtop transilluminator (Ultra-Violet Products, CA, USA) and amplicon sizes were determined by comparison to a 1 kb DNA ladder [20].

### Construction of *relA* mutants by Lambda Red Recombinase cloning

The Lambda Red Recombinase method as previously outlined [24] was used to construct *K. pneumoniae* ATCC 43816 KPPR1 and *K. pneumoniae* KP03 mutant strains. To make electrocompetent cells, *K. pneumoniae* electrocompetent cells which should contain pkD46 plasmid were made by growing strains overnight in LB broth containing ampicillin at 30°C with shaking. The following day, 2% (v/v) inoculum of overnight broth were inoculated into another fresh LB broth with ampicillin and allowed to grow at 30°C for 1 h, after which L-arabinose at 50 mM concentration was added and the broth cultures were further grown at 30°C until an optical density of 0.5 to 0.6 (approximately 3 h-exponential phase) at 600nm (OD_600_) was achieved. A 30 min incubation on ice was used for cells and then centrifuged in sterile cold 50 ml tubes at 8000 × g for 10 min at 4°C. Cells were serially washed after the supernatant was decanted, and further centrifuged in ice cold sterile 1 mM HEPES at pH 7.4 (Invitrogen, Carlsbad, California, USA), distilled water and 20 ml 10% glycerol. Pellets were stored at -80°C after resuspension in ice cold sterile 10% glycerol.

To construct *relA* mutants, *E. coli* BW25141 strains that contains the pkD4 plasmid (Coli Genetic Stock Centre) were extracted using 100 µl of DNA rehydration solution (TE buffer: 10mM Tris pH 8.0, 1mM EDTA) and stored at -20°C. Further, a 70-72 bp length oligonucleotide primers were designed using the Nucleotide BLAST program from NCBI by using a primer design template that contained 50 bp homologous to the target genomic locus to be deleted and 20 bp from the kanamycin cassette gene attached to the 3’ end. This kanamycin cassette is carried on the pkD4 plasmid which confers kanamycin resistance. PCR was used to generate the target linear fragment using the following parameters of 95°C for 5 min; 30 cycles of 95°C for 1 min, 58°C for 1 min, 72^0^C for 1 min and a final extension of 72°C for 5 min. The resulting amplicons from the PCR analysis were pooled and purified using the IBI Scientific PCR Purification kit and then digested overnight at 37^0^C with *Dpn*1 (New England Biolabs, Ipswich, MA, USA). Afterwards, 5 µl of the purified product was added to KP03/pkD46 and KPPR1/pkD46 electrocompetent cells and then incubated on ice for 10 min. A 1 mm cuvette was used to electroporate the mixture using an Eppendorf 1510 Electroporator; with field strength of 25kV/cm for 5.2 ms to treat cells. SOC medium was used to recover cells, and tubes were incubated at 30°C overnight with shaking at 180 rpm. Subsequently, cells were pelleted after incubation and resuspended in 25 µg/ml kanamycin/LB broth and the bacteria that were transformed were identified on kanamycin plates after incubation at 37°C overnight. Further, LB agar antibiotic plates were used to select for transformants after being restreaked and confirmation by colony PCR using flanking primers used for the creation of the mutants [6, 20].

### Complementation of mutants

Mutant strains were complemented by using the genomic region containing open reading frame of the *relA* gene which was PCR amplified, purified and cut with *Dpn*1 and then ligated with pGEM-T vector (Promega, Madison, Wisconsin, USA) after digestion with *Eco*R1 (New England Biolab [NEB]). The mutant *relA K. pneumoniae* strains were then complemented with the plasmid via electroporation. The presence of white colour colonies on LB amp/IPTG/X-Gal plates incubated after 18-24 h at 37^0^C and then placed in refrigerator at 4^0^C for further colony development was used to verify recombinant clones. The restoration of the deleted gene in mutant strains by the vector carrying the products of PCR were confirmed by using primers that flanked the desired gene using colony PCR [20].

### Environmental stress response assay

Osmotic, temperature and ethanol were used in this study for environmental stress assay. This is as a result of known information that microorganisms are constantly sensing and responding to environmental stimuli which include and not limited to osmotic stress, oxidative stress, starvation, changes in temperature and desiccation [20, 25]. In each stress assay, four clinical *K. pneumoniae* (KP03, KP04, KP18 and KP24) and *K. pneumoniae ATCC* 43816 KPPR1 strains (wild types) and two mutant (KP03*ΔrelA* and KPPR1*ΔrelA*) strains were used. Sensitivity to alcoholic and osmotic stress were measured by growing bacteria in LB broth (with kanamycin for mutants) at 37°C overnight. A 2% (v/v) culture was added to fresh LB broth after incubation and allowed to grow to mid log phase (OD_600_ of 0.4). Centrifugation of harvested cells followed and then M9 minimal broth was used to wash the cells twice, resuspensions in 0.9% saline to a cell density of 1×10^7^ CFU/ml. Diluted mutant and WT cultures were then added to M9 broth with 3 M of NaCl (osmotic) and 30% ethanol (alcohol) [25, 26], and incubated at 37°C. For every 30 min, periodic aliquots were taken followed by serial dilutions. Diluted cultures were then plated on LB plates at 37°C for 48h, and colonies counted. The same procedure above was used for the heat shock stress but no stressor was used. In addition, the diluted cultures were added to M9 broth media that was pre-warmed by placing flasks in a water bath at 67°C. Periodic aliquots were taken, and diluted cultures were plated directly onto LB agar plates to determine the viability of cells [26] and all experiments were done in triplicates.

### Measurement of nutrient starvation

To determine if wild-type *K. pneumoniae* strains (KP03, KP04, KP18, KP24, KPPR1) and *relA* mutants (KP03*ΔrelA*, KPPR1Δ*relA*) would be able to survive a 14 day starvation, cells were first grown in LB broth with antibiotics added to broth cultures that contained the mutant strains at 37°C overnight [20, 25, 27]. The cut-off period of 14 day incubation was used because of limited resources as well as the fact that WT strains were becoming too numerous to count after this time period, even though other studies have shown starvation up to 35 days for different species. Further, after incubation period, cells were allowed to grow to mid log phase (OD_600_ 0.4) at 37°C by adding a 2% inoculum to LB broth. Centrifugation at 10,000 × g for 5 min was used to harvest cells after growth and then washed twice in M9 broth minus nutrients followed by resuspension to a cell density of 1×10^7^ CFU/ml. The following medium were inoculated with cell suspension: medium I (carbon only), II (phosphate only), III (carbon and phosphate absent, IV (carbon and phosphate present) or V (carbon, phosphate and serine hydroxamate present). Cultures were incubated at 37°C at 150 rpm and for every 2-3 days periodic aliquots were taken, and cultures were done in triplicates and CFU determined.

### Young’s Capsule staining

Mutant and WT strains that were starved for 14 days along with non-starved culture strains were visualized for the presence and extent of polysaccharide capsule. This was done by using an inoculum from each culture, placing on a slide using a sterile loop, and then air dried afterwards. A 1% crystal violet (CV) solution was poured over the entire bacterial smears using a Pasteur pipette and allowed to incubate at room temperature for 7 min, and then 20% copper sulphate solution was used to wash off the CV solution. Before viewing under oil immersion, slides were then allowed to air dry for at least 20 min [20, 28]. To account for the average cell lengths of strains, approximately 100 individual bacterial cells from the different starvation media were counted.

### Biofilm and Agar Plate antibiotic susceptibility test

Mutant, WT starved, and non-starved cells were assessed for biofilm survival using 96-well polystyrene plates as previously described [29, 30] with some modification. Ceftazidime (CAZ) and/or gentamicin (GEN) antibiotic at a concentration of 50 µg/ml each was mixed with M9 minimal broth containing the starved and non-starved bacterial inoculum into a 96-well microtitre plate and then incubated at 37^0^C for 24h. An increased frequency of resistance to these two antimicrobial agents were shown in a previous study [20, 31], hence the reason for choosing them in this study. The removal of planktonic cells after incubation was done by washing twice in sterile water over a water tray, followed by staining with 0.1% crystal violet and then destaining using 200 µl 33% acetic acid for a final wash. A Victor microtitre plate reader (Perkin Elmer, USA) was used to measure the optical density of the destaining solution at 570 nm.

Non-starved, starved mutants and WT strains were used to carryout agar plate susceptibility assay. Non-starved mutant and WT strains were streaked on TSA agar plates and incubated at 37^0^C overnight. The following day, single colony were subsequently picked from these plates and placed in LB broth tubes and incubated at 37°C overnight. A 1:100 dilution of both starved and non-starved strains were used and a sterile glass spreader was used to spread 0.1 ml of the diluted cultures onto GEN and CAZ antibiotic plates. In addition, CFU was determined after plates (triplicates) were incubated at 37°C for 48h [17].

### Amplification of type 3 fimbriae (*mrkA*), phosphate (*phoR, phoU, pstB*) and sigma factor (*rpoD* and *rpoS*) gene

Chromosomal DNA was extracted from starved and non-starved (WT and mutant) strains using the Wizard Genomic DNA Purification Kit (Promega). Genes for type 3 fimbriae (*mrkA*), phosphate (*phoU, phoR, pstB* and sigma factors (*rpoS, rpoD*) were analysed using the primers identified in S1 Table 1. PCR mixtures consisted of 12.5μl 2x Promega Go *Taq* Green master mix reaction buffer (containing 1 U *Taq* DNA polymerase, 400 μM of each dNTPs, 3 mM MgCl_2_) and reaction buffers at optimal concentrations, ∼0.2 μg DNA, and 1.5μl (10 μM) of each primers (Integrated DNA Technologies, USA). Singleplex amplification parameters included a 5 min 94°C, followed by 35 cycles of denaturation at 94°C for 30 s, annealing at 62°C for 30 s and extension at 72°C for 2 min, and a final extension at 72°C for 10 min. All PCR components were added to pre-chilled tubes and were done on ice. Amplified PCR products were separated and visualized on ethidium bromide-stained 1.5% agarose, using Gel Doc UV illuminator (UVP BioDoc-it system).

### DNA sequencing

Biofilm, phosphate, sigma factor, *relA* and recombinant genes were sent to Eton Biosciences in San Diego, California, USA for sequencing. BLAST (Basic Local Alignment Search Tool) was used to compare the sequences obtained to known sequences in The GenBank database. DNASTAR Lasergene software by SeqMan NGen ®. Version 12.0. DNASTAR. Madison, WI, USA was used to edit sequences and SnapGene software from GSL Biotech; available at snapgene.com was used to view the chromatogram of sequences.

### Drosophila melanogaster infection feeding assay

The fruit fly infection assay was adapted from feeding assay developed by [32, 33, 34]. Single colony of mutant and WT strains were picked from freshly prepared streaked plate and placed in LB broth to grow overnight at 37^0^C. One percent of the inoculum was used to inoculate a fresh LB broth tube and cultures were allowed to reach mid log phase. For centrifugation, 1.5 ml of each culture was pelleted at 13000 x g for 5 min. The cells were resuspended in 170 μl of 5% sucrose solution. The resuspended pellets were spotted onto sterile filter disk (2.3 cm diameter Whatman paper) that was placed on the surface (5 ml of solidified 0.4% agar with 5% sucrose) at the bottom of a standard glass fly culture vial or in a 24-well plate. The vials were allowed to dry at room temperature for 30 min prior to addition of *Drosophila*. Adult female 3-5 day wild type fruit fly were starved for food and water for 5 h prior to being added to vials (10 flies per vial or 24-well plate). To anaesthetize the flies, they were placed on ice-cold tile throughout the sorting and transferring process. The vials were capped with cotton and then incubated at 25^0^C. The number of live flies to start the experiment were recorded and live flies were counted at 24 h interval over 14 days.

### Statistical analyses

The average CFU number corresponding to different treatment conditions were statistically analyzed using one-way analysis of variance (ANOVA) with Bonferroni post hoc test using PSPP version 4. A p value < 0.05 was considered significant.

## Results

### Construction of *K. pneumoniae ΔrelA* mutants and its complemented strain

The *relA* gene in strains KPPR1 and KP03 were mutated using the Lambda Red Recombinase method. A portion of the 200 bp (Fig 1A) fragment in the *relA* gene was deleted and replaced with a portion of a pkD4 plasmid that confers kanamycin resistance. Subsequently, agarose gel electrophoresis was used to verify the disruption of the *relA* gene by observance of a single band of 1550 bp that was seen for all four of the colonies used in colony PCR detection (Fig 1B). During the recombination of *K. pneumoniae*, growth of *ΔrelA* mutants were observed to be very slow in rich medium (approximately 48 hr for complete turbidity of flask using 4 colonies) compared to WT strains. The concentration of KP03 after 3 hr incubation was 3.33 x 10^6^ CFU/ml followed by KP03*ΔrelA* (6.03 x 10^3^ CFU/ml). As seen in photograph 1, KP03 had a large colony morphology and was mucoid and KPPR1 showed the characteristic hypermucoviscosity phenotype and larger colonies. However, complementation with the *relA* gene was unsuccessful after several attempts. possibly because the *relA* is tightly regulated within the cell and controls other components other than the stringent response. It could also be due to the growth defects in the mutant and secondary suppressor mutations might have accumulated.

**Figure 1A:**
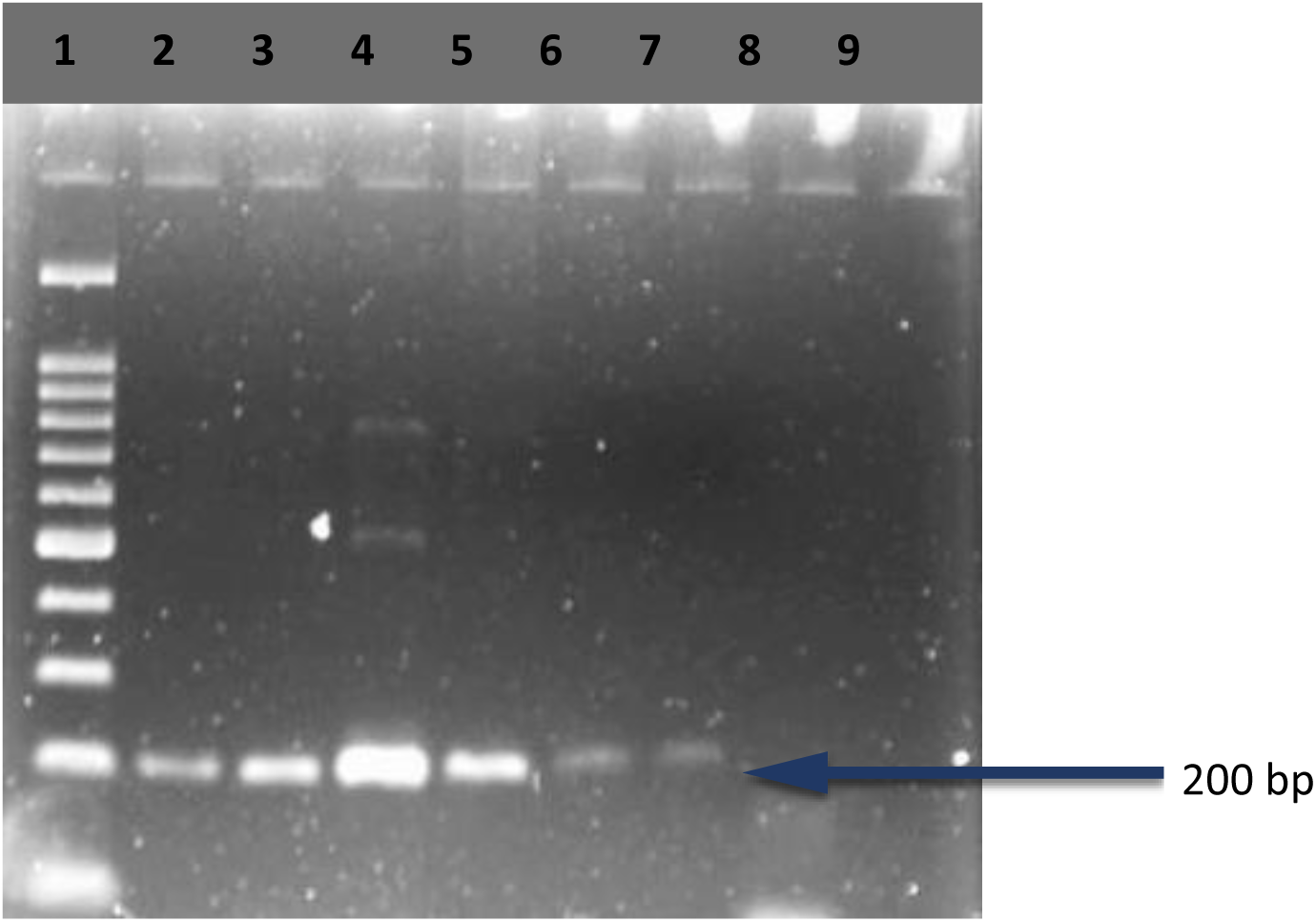
Electrophoretogram confirming the detection of *relA* gene. Lane 1, 100 bp ladder; lane 2, KP03; lane 3, KP04; lane 4, KP18; lane 5, KP24; lane 6-7, KPPR1

**Figure 1B.**
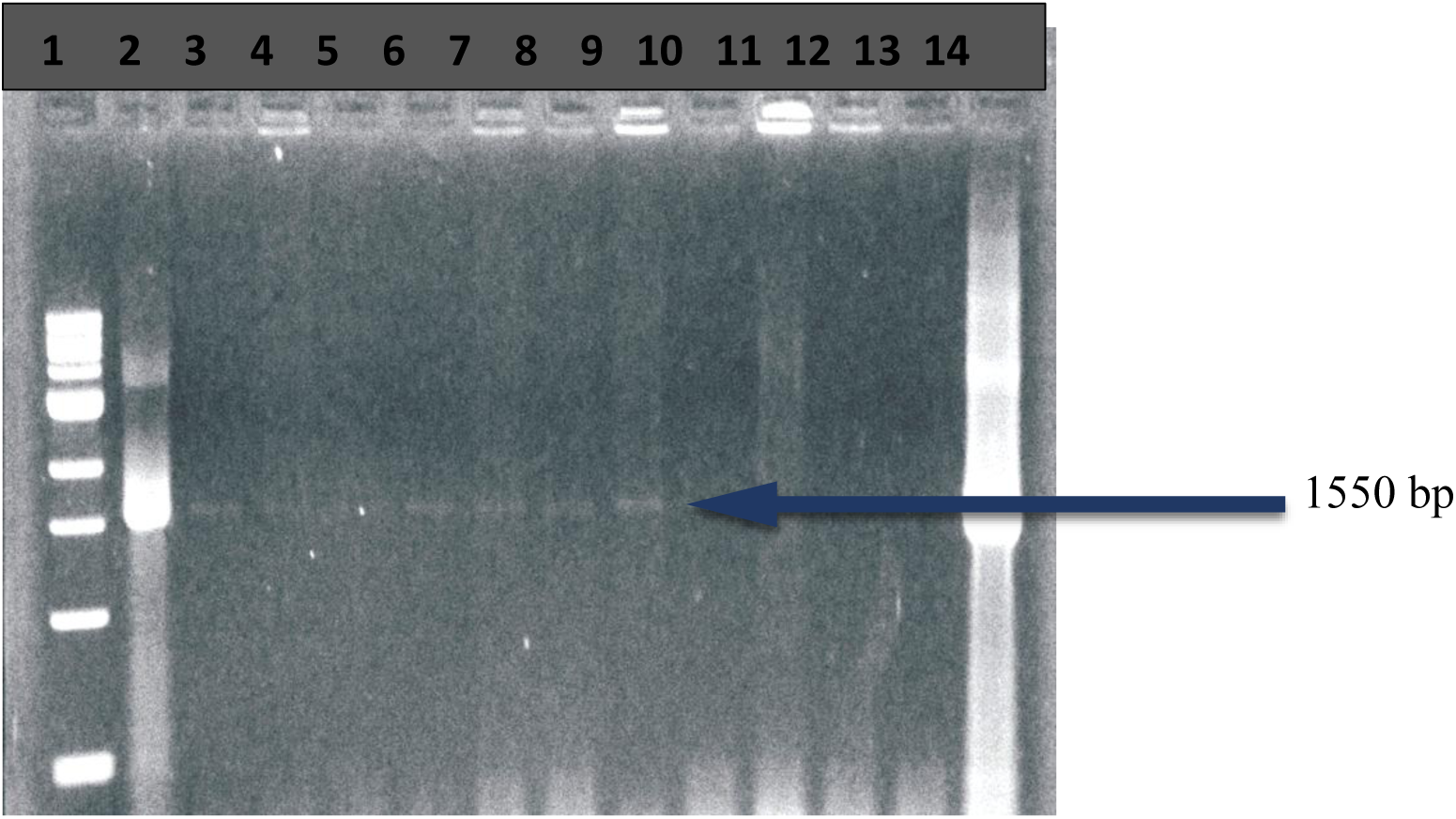
Electrophoretogram confirming the successful mutation of *relA* gene. Lane 1, MW marker; lane 2, linear kanrelA DNA; lanes 3-7, *KP03ΔrelA* colonies; lanes 8-9, *KPPR1ΔrelA* colonies; lanes 10-12, empty; lane 13, negative control; lane 14, linear kanrelA DNA

### *ΔrelA* mutant strains fails to survive under environmental stress

*K. pneumoniae* exponentially phased wild-types cultures (KP03, KP04, KP18, KP24, KPPR1) and *ΔrelA* mutants (KP03, KPPR1) were tested against environmental stresses of heat, NaCl (osmotic shock) and alcohol (ethanol) to determine if *relA* conferred any protection against the stressors. When WT and mutant strains were exposed to 30% ethanol, *ΔrelA* mutants were significantly more sensitive (0 log_10_ CFU/ml) to ethanol treatment over entire incubation period (120 min) in comparison to WT strains (Fig 3A). We also assayed if there was any protection mediated by *relA* to high temperature of 67^0^C and in the presence of high osmotic pressure (3 M). Data showed that the *ΔrelA* mutants were more sensitive (0 log_10_) to heat shock (Fig 3B) and osmotic stress over 120 min incubation than all the WT strains. Approximately 5-50% of the WT strains were able to cope with the heat stress at the 60 min incubation period (Fig 3C). One-way ANOVA showed statistically significant differences between *ΔrelA* and WT strains (p<0.05; 0.00), but no differences were observed between lengths of treatment for *ΔrelA* mutants (p>0.05; 1.00). Therefore, *relA* gene was required for maximum protection against heat, osmotic and ethanol stress.

**Figure 2:**
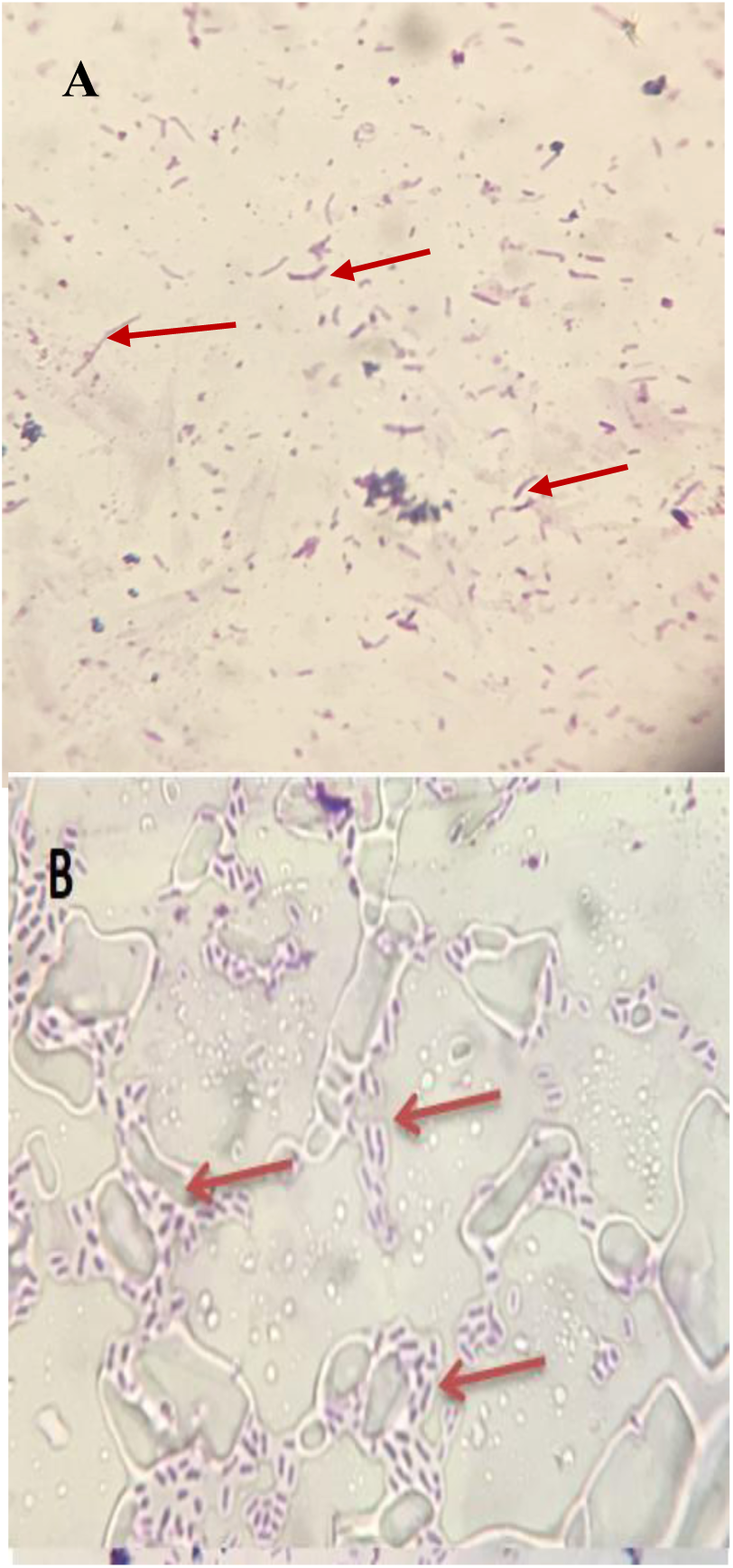
Photomicrographs of (A) KPPR1*ΔrelA* mutant from media without carbon, showing absence of capsule and presence of vegetative cells with elongated rods (arrows) (B) KPPR1 WT strain in non-starved media showing presence of capsule with clear zone around vegetative cell stained purple by crystal violet (arrows); Viewed under oil immersion.

**Figure 3.**
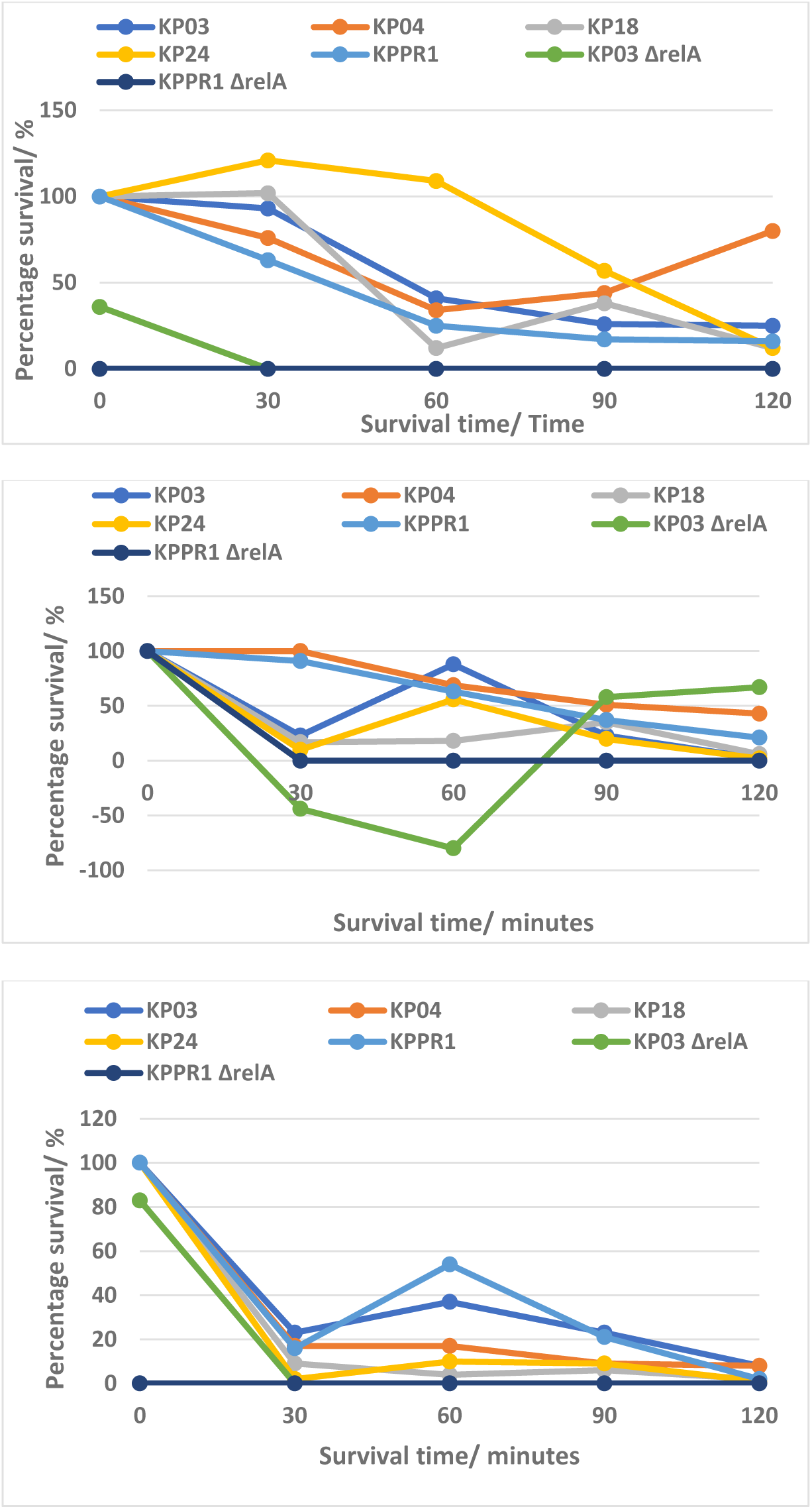
Percent viability of exponential-phase wild-type and *relA* mutant *K pneumoniae* strains in M9 media with (A) 30% ethanol; (B) 3 M sodium chloride; (C) at 67°C. Data represent the means of triplicate values relative to untreated strains.

### *ΔrelA* mutant strains fails to survive under nutritional starvation

Mutant and WT strains were subjected to prolonged carbon, phosphate, amino acid and elimination of carbon/phosphate starvation in order to determine the effect of gene mutation on the strains survival in relation to the general stress response system. This was further monitored by plating on either medium I, II, III, IV or V with 0.02% glucose in medium I to minimize the possibility of substrate accelerated death. Firstly, under carbon starvation (medium I), KPPR1*ΔrelA* mutants showed no growth and KP03*ΔrelA* had a high log value which indicated that both *ΔrelA* mutants showed a significantly increased sensitivity to prolonged carbon starvation when compared to WT strains. While under phosphate starvation (medium II), all WT strains showed a decreased log_10_ (CFU/ml) growth during the first 5 days, and then no growth was observed from 9-14d. However, no growth was observed for both *ΔrelA* mutants over 14 day incubation. One-way ANOVA showed there were significant difference between survival times for all WT strains (p>0.05; 0.00) and KP03*ΔrelA* (p<0.05; 0.00), but none was observed for KPPR1*ΔrelA* (p>0.05; 0.47) (S2 Table).

In addition, the elimination of carbon and phosphate from media (III), showed that WT strains log_10_ value ranged from 2.08 to 3.04 over 5-day period but there was a subsequent drop in viability of strains to 0 log_10_ from 7-14d. However, no value was determined for both *ΔrelA* mutants because strains were unable to grow in media lacking both nutritional requirements. Therefore, the data suggested that *ΔrelA* mutants showed an increased sensitivity to prolonged starvation within this media compared to WT strains. One-way ANOVA analysis showed significant differences between the treatment time for all WT (p<0.05; 0.00) (S2 Table).

Furthermore, to induce amino acid starvation in the strains, serine hydroxamate (SHX) was added (medium IV). There was a decreased sensitivity to treatment in KP03*ΔrelA* mutant {higher (positive) log_10_} but KPPR1*ΔrelA* mutant had no log value because no growth was observed, while all WT strains showed a negative log_10_ value that increased from -6.00 to -8.99 from days 2-12. A negative log value for WT starved strains meant that they were performing significantly less better than their non-starved and *ΔrelA* mutant counterparts during the course of the treatment and it also indicated that the mutant strains were not greatly affected by the addition of SHX, which resulted in a decreased sensitivity. Statistically significant differences were observed using between the treatment time for KP03, KP18, KPPR1 and KP24 (p< 0.05; 0.00, 0.01) but no difference was seen between survival times for strain KP04 (p>0.05; 0.09) and KP03*ΔrelA* (p< 0.05; 0.00) (S2 Table).

### Deletion of *relA* gene affects biofilm formation/survival of *ΔrelA and* WT strains to gentamicin and ceftazidime

WT and mutant strains were assessed for the extent of biofilm survival before and after starvation when treated with GEN and CAZ (Fig 5A-5E). With the exception for KP18 which had greater than 100% survival before starvation for all five treatment conditions, all other WT strains had approximately 30-50% survival [20] and *ΔrelA* mutants showed 64-109% survival.

**Figure 4.**
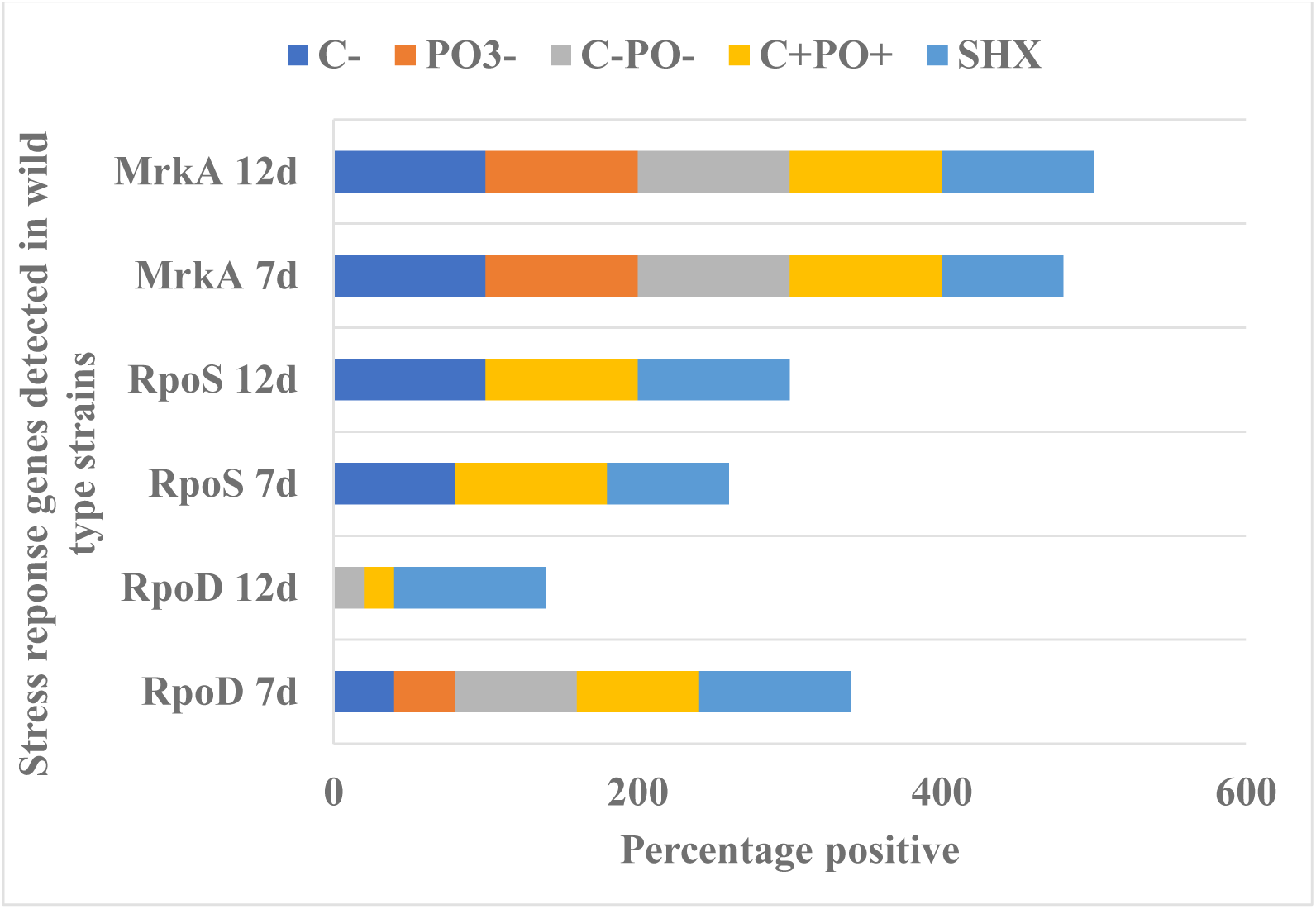
*K. pneumoniae* WT and mutant strains showing percentage positive detection of stress response genes during nutrient starvation after 7 day and 12 DNA extraction

**Figure 5.**
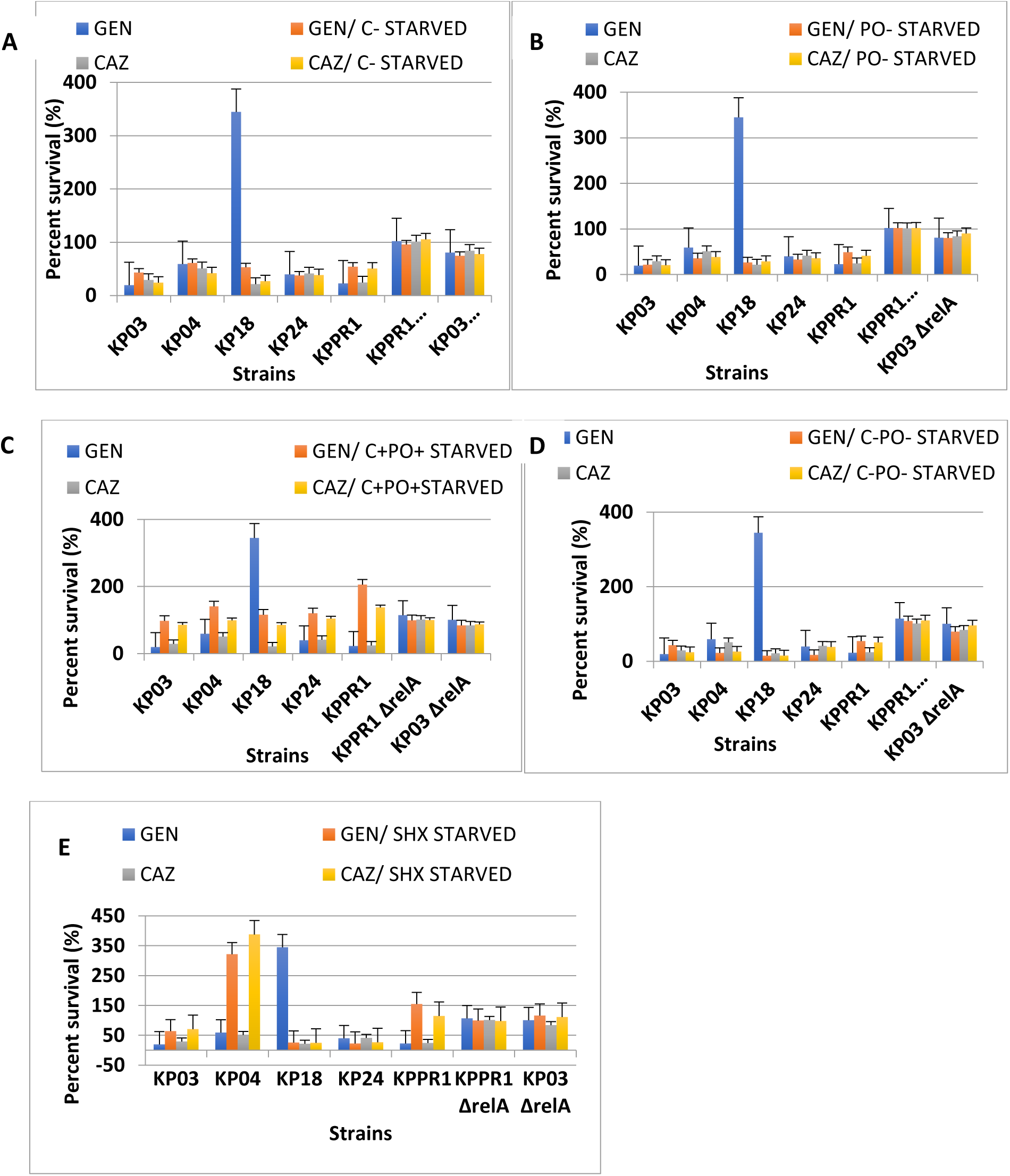
Percent survival of wild type and *relA* mutant *K pneumoniae* biofilm cells during gentamicin and ceftazidime treatment after 14 days of starvation (A) without carbon present; (B) without phosphate present; (C) with carbon and phosphate present; (D) without carbon and phosphate present; (E) with amino acid starvation. Values represent the means of experiments performed in triplicate.

After carbon starvation with GEN present as illustrated in Fig 5A, there were higher biofilm survival for *ΔrelA* (75-96%) mutant strains compared to WTs (38-61%) (Fig 5B) and a similar trend was observed after phosphate starvation *ΔrelA* (80-102%) mutant strains; WTs (20-49%)). On the other hand, after carbon and phosphate present starvation with GEN added (Fig 5C), biofilms from *ΔrelA* (84-99%) mutants were slightly more sensitive to GEN than WT (98-206%) strains. Carbon/phosphate minus after starvation with GEN biofilm survival (Fig 5D), showed that *ΔrelA* mutants were more resistant than biofilms of WT strains (15-54%). Furthermore, after amino acid starvation with GEN present (Fig 5E), the percent biofilm survival of WT strains KP04 and KPPR1 showed the highest biofilm growth (respectively) followed by KPPR1*ΔrelA* (99%) and KP03*ΔrelA* (116%) and the other three WTs. There was no statistically difference were observed based on one-way ANOVA (p< 0.05; 0.00, 0.01)

Considering exposure of strains to CAZ antibiotic, approximately 22-41% biofilm survival rate was observed for all WT strains for the five treatments before starvation and this is represented in Fig 5A-5E. Also, on Fig 5A after carbon starvation with CAZ present, biofilm survival of *ΔrelA* (78-106%) were higher than WTs (24-42%) strains and a similar trend was observed after phosphate starvation where *ΔrelA* biofilm survival was 90-102% and WT strains were 21-41% (Fig 5B). Furthermore, after carbon and phosphate present starvation with CAZ present, WT strains had a slightly high biofilm survival rate (86-135%) than *ΔrelA* (87-100%) (Fig 5C) but after carbon/phosphate minus with CAZ present biofilm WT strains were more sensitive to antibiotic (15-51%) than *ΔrelA* mutants (98-110%) (Fig 5D). On the other hand, the presence of CAZ in media after amino acid starvation showed that KP03*ΔrelA* (111%) and KPPR1*ΔrelA* (98%) biofilms were more resistant than WT strains except for KP04 (388%) as seen in Fig 5E. One-way ANOVA showed that statistically significant differences (p< 0.05; 0.00) were observed between groups of treatment with non-starved and starved WT strains as well as starved *ΔrelA* mutant strains on exposure to CAZ.

### *ΔrelA* mutants’ planktonic cells fails to survive under gentamicin and ceftazidime antibiotic stress

Agar plate antibiotic susceptibility assay was used to test mutants’ and WT strains’ ability to cope with antibiotic stress. According to Fig 6A which represents strains plated on CAZ agar plates after starvation treatments, *ΔrelA* mutants (0-1%) and WT KPPR1 (2%) carbon-starved strains were more sensitive to CAZ than the other WT strains. Among phosphate starved and carbon/phosphate minus-starved and non-starved strains (LB/CAZ) strains, this trend was also observed. However, after amino acid starvation, KP24 (783%) and KPPR1 (357%) were significantly more resistant to CAZ than all other strains followed by KP03*ΔrelA* (100%). One-way ANOVA showed statistically significant difference between groups of treatment (p< 0.05; 0.00) for all strains. Bonferroni post hoc test showed statistically significant differences were observed for non-starved strains (LB/CAZ) versus carbon-starved, carbon/phosphate-starved, phosphate-starved, and amino acid-starved counterparts.

**Figure 6.**
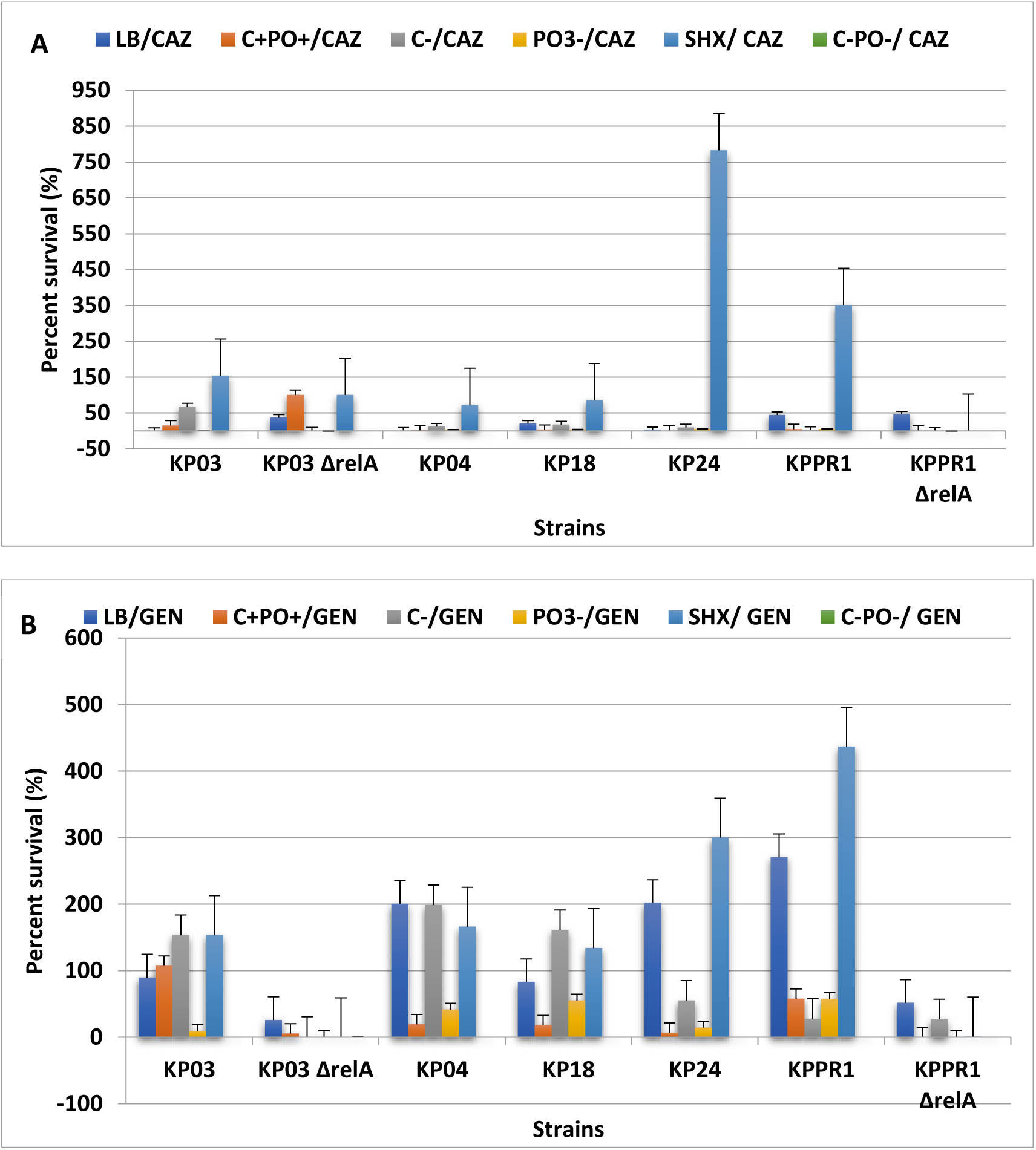
Percent viability of *K. pneumoniae* strains during (A) ceftazidime (CAZ) or (B) gentamicin treatment after 14 days of starvation and non-starvation. Data represent the CFU means of triplicate values.

Conversely, GEN agar was used to grow non-starved WT strains and as seen in Fig 6B, they performed better under this antibiotic treatment than *ΔrelA* mutants. After carbon starvation, KP03Δ*relA mutant* (1%) was significantly more sensitive than WT strains (KP03, KP04, KP18). Among carbon/phosphate present-starved and carbon/phosphate minus-starved strains, this trend was also observed but the reverse was observed for phosphate-starved strains. Furthermore, amino acid starved Δ*relA* mutants showed no growth; but all WT strains showed decreased sensitivity to GEN especially KPPR1 (437%). No statistically significant difference between groups of treatment (p > 0.05; 0.39) for all strains using one-way ANOVA. However, Bonferroni post hoc test, showed significant differences with strains KP03, KP04 and KP18, and carbon-starved bacteria exposed to GEN was more resistant than phosphate-starved, carbon-/phosphate minus-starved cells, and even non-starved cells (LB/CAZ).

### Mutations in *relA* of *K. pneumoniae* results in absence of capsule

All non-starved WT strains with carbon and phosphate present in the media showed the presence of capsule and all starved WT strains with both nutrients present showed the absence of a capsule. However, for carbon starvation, three of the strains showed capsule present (KP03, KP04, KP18) (Appendix D.1) but under phosphate starvation, only KP04 had capsule and none detected in the other WT strains (Appendix D.2). In addition, no capsule was present for the other three-starvation treatment. Among *ΔrelA* mutant starved strains, capsule was absent, and all non-starved mutants had no capsule present as well. Based on phenotypic observations, mutant untreated strains formed small colonies compared to WT untreated strains on LB agar plates, and were less mucoid, suggesting there might be a defect in capsule biosynthesis. This was also observed for some mutant strains under carbon and phosphate starvation.

### Mutations in *relA* of *K. pneumoniae* resulted in altered cell length

When KPPR1Δ*relA* mutant strain was grown under carbon starvation, the average cell length was considerably longer than WT strains, showing size of approximately 3.58 μm, whereas the average length of KP03*ΔrelA* was slightly longer than WT strains, reaching length of 1.92 μm. In contrast, the average length of KP03*ΔrelA* mutant cells when both nutrients were included in growth medium was 2.56 μm which is much longer than that of WT strains under same condition. Furthermore, KP03*ΔrelA* had considerably shorter rods than WT strains under amino acid starvation, reaching average length of 1.14 μm (Fig 2A-2B). However, the length of the majority of KP03*ΔrelA* under phosphate starvation were the same as with all other WT strains.

### Majority of stress response genes undetectable in *ΔrelA* mutants

The amplification of stress response genes in *K. pneumoniae* were determined by PCR analysis of non-starved WT and *ΔrelA* strains after 7 and 12d starvation and are represented in Fig 4. There was no detection of phosphate genes (*phoU, phoR* and *pstB*) in mutants and wild-type strains after 7d and 12d starvation as well as non-starved strains. All non-starved WTs and mutant strains showed the presence of *rpoD* and *mrkA* genes, but there was no detection of *rpoS* gene. All WT strains on both days were positive for *mrkA* gene while *mrkA* gene was not detected in KP03*ΔrelA* but was detected in phosphate, amino acid and carbon phosphate present starvation for KPPR1*ΔrelA.* In addition, *rpoD* gene was only detected on day 7 for all WT strains and *rpoS* gene was not detected in KP03*ΔrelA* strain over both days but was detected in KPPR1*ΔrelA* during carbon-, phosphate and amino acid starvation.

### *Drosophila melanogaster* infected with wild-type and mutant strains of *K. pneumoniae* through feeding assay

Single mutants of *ΔrelA* strains versus wild-type strains of *K. pneumoniae* were assessed for their ability to kill *Drosophila,* and fly survival was monitored over 14 days. The uninfected control consisted of inoculated filter disk with 5% sucrose solution. Based on Kaplan-Meier survival curves illustrated in Fig 7, the uninfected control only had one fly dying over the 14 days incubation which was observed at 9d. Wild-type strain KP18 were fairly harmless slowly killing only a minority of the flies over the first 5 days. On the other hand, KPPR1*ΔrelA* and KP03*ΔrelA* strains were highly effective in killing flies compared to WT strains. In addition, KP03 was the only WT strain with a significant increase in fly kill and then a steady death rate from 2-6 days but then decreased in survival after 7 days. Both *ΔrelA* mutants were capable of killing flies over the course of incubation but with reduced effectiveness except for KPPR1*ΔrelA* mutant strains when compared to WT strains. However, there was no significant difference in killing among the *ΔrelA* mutants and the WT strains (p>0.05; 1.00) using one-way ANOVA.

**Figure 7.**
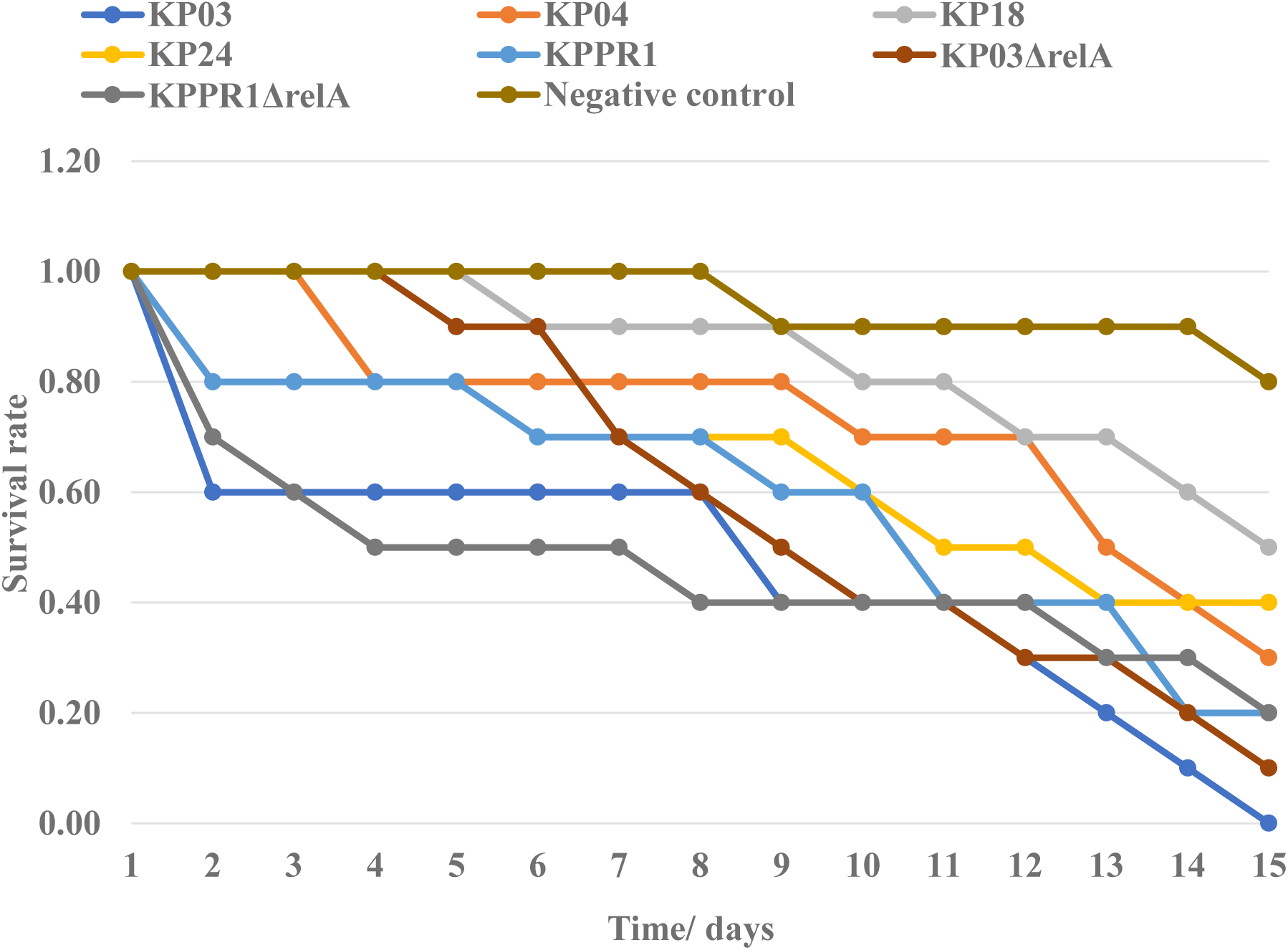
Virulence of the *K. pneumoniae* mutants (*ΔrelA*) strains in fruit flies. Graph represent the survival rate (death) of flies over time. The values are the averages from three replicate experiments over, each with 10 flies, and the SEM is shown for each value plotted

### Sequencing of stress response genes

A total of 16 PCR amplified DNA samples from KP03 and KPPR1 starved and non-starved strains were sent for sequencing to Eton Biosciences in San Diego, California, USA. However, only 10 of those samples were able to be analysed using the DNASTAR and SnapGene software. Majority of the amplified gene products from the different strains used in this study were similar to other *K. pneumoniae* species found in the Genbank database. There was also similarity with another *Enterobacteriaceae*, *E. coli* and *Providencia* sp. There was no difference in sequence of starved and non-starved strains based on those samples that could be analysed.

## Discussion

Bacterial stress inflicts a variety of adaptive and protective responses. Most bacteria live in a dynamic environment where temperature, availability of nutrients and presence of various chemicals impact its survival [35]. Based on the findings in this study, it was observed among the different environmental stress treatments: the longer the strains remained in the stress medium, the less viable they became. However, there was a significant reduction in cell viability for KP03*ΔrelA* and KPPR1*ΔrelA* mutants over all environmental stress tests. This same trend was observed in *ΔspoT* mutants of *K. pneumoniae* [20]. Therefore, the presence of the *relA* locus was important for the survival of *K. pneumoniae*.

It is evident from the results obtained that under phosphate starvation; WT strains performed much better than under glucose starvation [20]. While phosphate is required for biosynthesis of critical cellular components such as ATP, protein synthesis, nucleic acid and phospholipids [36, 37] a carbon (energy) source was more important. These results were expected but it doesn’t negate the fact that a media void of phosphate showed significant differences in survival of strains. Obviously, the survival of the strains could be influenced by other factors not limited to but could include the possibility of carbon and phosphate sensing genes expressed during starvation [22].

Comparison of WT strains with Δ*relA* mutants under carbon-, phosphate- and amino acid starvation showed that *ΔrelA* mutants performed significantly less favorable than WT strains and it is an indication that replication and synthesis of certain genes required for growth were stalling in mutant strains. This does not corroborate results observed in *ΔspoT* mutants of *K. pneumoniae* which had increased survival in all starvation treatments [20]. In addition, in *ΔrelA* mutants, it was noted that disruption of this gene was linked to amino acid starvation and thus strains were less viable under such conditions, and the presence of the remaining *spoT* gene was not enough to overcome the stress induced under this condition and the other starvation conditions. Synthesis of the alarmone ppGpp is usually expected with the presence GTP and ATP in cells, and often times expression of stress response processes is elevated, and a reduction ensues in ‘housekeeping’ processes [39, 40]. Therefore, it could be inferred that the stringent response was induced in media lacking both nutrients, but it was significantly observed in phosphate starved medium. In addition, the absence of the *relA* gene resulted in an impaired stringent response system and consequently resulted in strains being less able to survive and as such less virulent.

Investigation of biofilm formation and starvation-survival responses in varying concentration of antibiotics were studied. Data showed that within antibiotic pressure, starved WT strains continued to be resistant to antibiotics with observations markedly seen in carbon-starved cells than phosphate-starved cells. In addition, with the presence of carbon and phosphate in the media, a 206% survival rate was observed for WT starved, and non-starved strains were more tolerant to gentamicin than ceftazidime exposure. This could be as a result of the low oxygen level found in biofilms to which gentamicin has a weak penetrative effect towards. Furthermore, a 20-50% survival rate was observed for strains exposed to ceftazidime when compared to non-exposed strains [20]. The presence of resistance genes SHV and CTX-M that were already detected in strains should not have given such a result since they confer extended spectrum β-lactamase resistance phenotype [41, 42].

Over the years there have been controversy among the scientific community surrounding the role the stringent response plays in biofilm formation. Age of biofilm, environmental conditions for growth and by extension the flow rate could be possible reasons [43]. On the other hand, biofilm production in the *relA* mutants following starvation were unexpectedly enhanced in the presence or absence of antibiotic pressure, and under amino acid starvation, the biofilms of mutant strains performed better than that of the starved WT strains. A similar observation was observed for *ΔspoT K. pneumoniae* mutants [20]. Hence, *relA* mutation contributes to antibiotic tolerance in biofilm, and according to two studies done on *relA* mutation and its effect on biofilm formation and antibiotic tolerance, it was hypothesised that this tolerance could be as a result of resting alarmone basal levels which may confer antibiotic tolerance of significant proportion in immunocompromised situations [44, 45, 46]. To further hypothesize that the presence of an intact *spoT* gene which is known to have a weak hydrolytic activity of (p)ppGpp could have contributed to this basal level concentration of the alarmone and render conventional antibiotic treatment recalcitrant. Several previous studies have shown that the lack of (p)ppGpp resulted in decreased biofilm formation in *Enterococcus faecalis* [57], *Vibrio cholerae* [58] *Bordetella pertussis* [59]. However, work done on *Actinobacillus pleuropneumoniae* S8 showed that (p)ppGpp deletion mutant *ΔrelA/spoT* of the bacteria produced significantly more biofilms than WT and complement strains [60]. This was also seen in other bacteria such as *Francisella novicida* [61] *Pseudomonas putida* KT2440 [62] and *Porphyromonas gingivalis* [63]. Indeed, these results contrasts the previous studies for reasons that are unclear because the mechanisms involved have not be explored but does mirror this present research on *K. pneumoniae*. In addition, detection of the biofilm gene *mrkA* which codes for type 3 fimbriae, mediates attachment to- and biofilm formation on abiotic surfaces in-vitro, was an important finding. This gene was found present in all starved and non-starved WT and KPPR1*ΔrelA* starved mutants but was not detected in KP03*ΔrelA* starved mutants. The latter should not be interpreted that the strain could not form functional biofilm structures because research has shown that bacteria that are fimbriated, that possesses no functional adhesin can still facilitate biofilm formation on abiotic surfaces [47].

It is known knowledge that antibiotic tolerance is one of the most important phenotypic features linked to (p)ppGpp [64]. Hence, the effect of *relA* deletion on antibiotic susceptibility was done using the agar plate susceptibility assay. The results showed that *ΔrelA* mutants exposed to gentamicin and ceftazidime showed increased susceptibility when compared to WT strains. Similar work done on *ΔspoT K. pneumoniae* mutants showed an increased antibiotic susceptibility to GEN and CAZ [20]. This is in contrast to work done on *Mycobacterium smegmatis.* Data from this study showed that *Δrel* (mutated from *rel_msm_* which has bifunctional properties) had a higher tolerance to most antibiotics [64] than *ΔrelZ* (mutated from *relZ* which is short alarmone synthetase that can make and not hydrolyse (ppGpp)) displayed sensitivity to most antibiotics [65]. Therefore, the authors hypothesized that the influx/efflux that is created by the antibiotics affected these strains due to the altered (p)ppGpp homeostasis. Consequently, this accumulation of ppGpp would likely inhibit peptidoglycan synthesis as well binding to the 30S ribosome causing misreading of t-RNA and inability to synthesis proteins that were required for growth [35].

*K. pneumoniae* usually produces a polysaccharide capsule which is a major virulence factor in the bacteria. The absence of a carbon source such as glucose should have resulted in absence of capsule surrounding the strains. However, capsule was detected in the three WT carbon-starved strains of KP03, KP04, KP18 [20]. The formation of capsular structures does not require glucose hence it was detected in the strains mentioned before, but it is known to play an important role the architecture of the capsule [48]. During antibiotic exposure, there was a statistically significant difference in viability of strains with capsule versus those strains without capsule, with the exception of KP04, and there was no polysaccharide capsule present for phosphate-starved cells [20]. Therefore, it seems that it is the presence of phosphate which helps in the formation and assembly of phosphorylated sugars and other carbohydrates are needed for the synthesis of the capsule [49]. Furthermore, when carbon and phosphate were present in the media, non-starved WT strains produced a capsule while starved WT strains produced none. Interestingly, the survival of *relA* mutants proved to be linked to the presence of polysaccharide capsule because under all five starvation conditions, both *relA* mutants saw the loss of the capsule. Also, based on the data, specific nutrients were required by strains for synthesis of capsule and components of the stringent response, especially the *relA* gene. Hence, there is a possibility that a defect in capsular synthesis resulted from a mutation in *relA* gene, thus resulting in a decrease in viability of mutant strains even upon environmental stress exposure and especially under antibiotic stress.

Starved and non-starved WT cell had similar cell length despite the different treatment conditions. However, KPPR1*ΔrelA* mutant cells were much longer than KP03*ΔrelA* and WT cells in all starvation media and KP03*ΔrelA* on the other hand had the second longest averaged cell length, greater than all WT cells. These findings suggested that a reduction in *K. pneumoniae* cell lengths of WT cells and an increased cell length of *ΔrelA* mutants might be as a result of the accumulation of the alarmone in these cells. Also, work done on *Mycobacterium smegmatis* showed that a lack of a SR altered cell morphology- cells appeared significantly longer for mutants than WT cells [66]. In addition, research have shown a link between short cell lengths and the capacity of these strains being more antibiotic resistant and even increased resistant to oxidative and pH stress which eventually causes an increase persistent survival under extreme environmental conditions [50, 51].

Within *E. coli* and *P. aeruginosa*, phosphate metabolism and management (including detection and responding to environmental phosphate concentration changes) are controlled by the Pho regulon. This Pho regulon is a two component PhoB-PhoR system that detects low inorganic phosphate level in cells [38]. Our assumption was that regulatory genes could play a role in survival of strains under phosphate starvation but based on PCR analysis, *phoU, phoR* and *pstB* gene were not detected in non-starved, starved WT and mutant strains even though genes are conserved within the bacteria’s chromosome. Therefore, lack of amplification might be as a result of contamination of DNA with an inhibitor in the case of non-starved strains, since concentration of template DNA was sufficient for high yields, but low DNA template concentration and/or inhibitor could account for lack of amplification in starved strains. Also, *rpoS* and *rpoD* was also undetectable in both *ΔrelA* mutants on both DNA extraction days. Therefore, the lack of amplification of *rpoS* and *rpoD* could be due to concentration of DNA template for amplification. For instance, extraction of DNA from starvation media for the different strains had low purity, especially observed in phosphate and combined phosphate media and in some instances the ability to form pellet was also absent.

NCBI BLAST program was used to compare 10 nucelotide sequences from *K. pneumoniae* strains used in this study with a database of sequences in the library in order to identify organisms similar to them. *K*. *pneumoniae* strain I72 was the only strain which showed a consistent similarity with PCR amplified genes of *relA, mrkA, rpoS, rpoD* followed by *E. coli* isolate EC-12536 (*mrkA and relA*). However, there was no published data on the strain from the authors that submitted them to the Genbank in late 2018. Other similarities were seen for *rpoD* and *rpoS* gene with *K. pneumoniae* strain ST23 and KP14. ST23 is a carbapenemase-producing hypervirulent *K. pneumoniae* isolate encoding blaKPC-2 plasmid, *rmpA, rmpA2 iroBCDN, peg-344* and *iucABCD-iutA* genes [52]. KPPR1 *mrkA* gene showed 99.41% similarity to *P. rustigianni* but the prevalence of published information on this species is low and the most popular species from the genus are *P. stuartii* and *P. rettgeri.* General characteristics of this genus is that they are opportunistic pathogens of humans, members of the *Enterobacteriaceae* family and common cause of UTI infections [53]. In addition, *P. stuartii* causes 9% of infection in patients with long term hospitalization [54] and is shown to be good biofilm-producers [55].

In the fruit fly, *D. melanogaster* model, adult female fruit flies were fed on lawns of *K. pneumoniae* WT and *ΔrelA* and mutant strains over a 14 day period. *D. melanogaster* was used because of its similar innate immune system to humans that *K. pneumoniae* usually infects and cause disease in these immunocompromised individuals. In this study it was shown that KPPR1 strains with *relA* single deletion had an increased fly kill than the other strains but was still deemed not significant. Two hypotheses have been proposed that could have contributed to the increase death of flies with mutants. Firstly, differences in killing between mutants and WT strains could be as a result of the gut microbiome of the fruit fly. Mutations usually results in a fitness cost on a bacterium and hence in this situation mutants would outcompete the microorganisms in the GI tract thus killing flies faster than WT strains that had no mutations and hence, killed flies more slowly. Secondly, single mutants of KP03 and KPPR1 is hypothesised to be able to produce basal levels of the alarmone. Therefore, this basal level of ppGpp production could also constitute one of the key factors in controlling virulence and pathogenesis in *D. melanogaster* models than an intact stringent response system. Even though data showed there is no statistically significant difference, it revealed that an intact stringent response is not required for virulence of the bacteria. Even if there is one defective gene and the other is fully functional, as long as small amount of the alarmone is able to produce, a coordinated response can still be facilitated which could aid pathogenicity and impaired treatment regimen. Also, a study done showed that *Δrel and Δrshb* single mutants of *P. gingivalis* 381 killed *Gallaria mellonella* at comparable rates to parent strains at approximately 75% after 78 hr of infection. This meant that the signalling of (p)ppGpp impacted the virulence of *P. gingivalis* strain 381 [63].

## Conclusion

When *K. pneumoniae* underwent nutritional and environmental stress, the data showed that the *relA* gene was pivotal in its survival and proliferation throughout treatment. Research has suggested that cells in exponential phase of growth possesses complex machineries that help bacteria to survive upon exposure to diverse stressful conditions. Hence, the findings in this report suggested that starvation and environmental stress aided *K. pneumoniae* in the loss of its main virulence factor, the capsule, especially when phosphate was eliminated from the media and along with the disruption of the *relA* locus. The biofilm formation gene *mrkA* could account for increased biofilm formation for mutants, but it would appear that (p)ppGpp (*relA*) negatively regulates biofilm formation in *K. pneumoniae* as observed in *F. novicida*, *A. pleuropneumoniae* S8 and *P. putida* KT2440, and mutants were more vulnerable to antibiotic pressure in agar plate susceptibility test. Overall, sequenced stress response genes used in this study showed high similarity to other *Enterobacteriaceae* in the Genbank database which could mean that amplified genes are conserved sequences among the family. In addition, by using the *D. melanogaster* model for pathogenicity, we were able to demonstrate that the stringent response does not require *relA* for eliciting full virulence in fruit flies but the presence of a either one of the gene which is fully functional can still have adverse effect on insect survival. Even though a consistent direct relationship was not observed between single mutants of *relA* and the various phenotypes seen in this study, differences might be explained via differences in the intracellular ppGpp concentration levels which determine the exact gene expressions controlling the phenotypes in different conditions including environment, nutritional and antibiotic stress [56]. Taken together, we conclude that the *relA* gene was an important RSH component, since the stringent response was inhibited by its absence. This was observed in *△relA* mutants which were more susceptible to antimicrobial, environmental and nutritional stress, absence of capsule, longer cell length and increased biofilm formation when compared to WT strains.

## Acknowledgments

Special thanks to the staff of the University Hospital of the West Indies and Central Medical Laboratory, Kingston, Jamaica, for assisting with initial strain collection, and the *E. coli* Genetic Stock Center (CGSC) for providing the pkD46 and pkD4 plasmid.

